# Comparable environmental stability and disinfection profiles of the currently circulating SARS-CoV-2 variants of concern B.1.1.7 and B.1.351

**DOI:** 10.1101/2021.04.07.438820

**Authors:** Toni Luise Meister, Jil Fortmann, Daniel Todt, Natalie Heinen, Alfred Ludwig, Yannick Brueggemann, Carina Elsner, Ulf Dittmer, Stephanie Pfaender, Eike Steinmann

## Abstract

The emergence of novel SARS-CoV-2 B.1.1.7 and B.1.351 variants of concern with increased transmission dynamics has raised questions regarding stability and disinfection of these viruses. In this study, we analyzed surface stability and disinfection of the currently circulating SARS-CoV-2 variants B.1.1.7 and B.1.351 compared to the wildtype. Treatment with heat, soap and ethanol revealed similar inactivation profiles indicative of a comparable susceptibility towards disinfection. Furthermore, we observed comparable surface stability on steel, silver, copper and face masks. Overall, our data support the application of currently recommended hygiene concepts to minimize the risk of B.1.1.7 and B.1.351 transmission.

## Background

Since the outbreak of *Severe Acute respiratory Syndrome Coronavirus-2* (SARS-CoV-2) at the end of 2019, > 120 million cases and > 2.8 million death (March 31^st^ 2021) have been reported [1]. Viral evolution includes the natural emergence of viral variants, which can encode for a variety of mutations in their genome compared to the parental wildtype virus. Mutations which confer either enhanced fitness, higher pathogenicity, better transmissibility or immune escape are of special concern as they could significantly influence transmission dynamics with devastating consequences. Independent lineages of SARS-CoV-2 have recently been reported: UK, B.1.1.7; South Africa, B.1.351; and Brazil, P.1 [2]. Importantly, these variants of concern (VOC) display higher reproduction numbers than preexisting variants and consequently increase incidences in various countries. Moreover, VOCs have been associated with more severe course of infection and/or potential immune escape due to multiple changes in the immunodominant spike protein [3–5]. Since the global access to COVID-19 vaccines is still limited, diligent attention on transmission-based precautions is essential to limit VOC spread. However, given the rapid spread and increased transmission dynamics of the emerging variants, concerns regarding the effectiveness of current hygiene measures and inactivation strategies have been raised. Here we compared the stability of three SARS-CoV-2 strains, the preexisting B1.1.70 variant (herein referred as WT virus) and the currently emerging B.1.1.7 and B.1.351 variants on different surfaces and their sensitivity to heat, soap and ethanol.

## Methods

### Viral isolates and Cell culture

For SARS-CoV-2 virus suspension preparation, Vero E6 cells (kindly provided by C. Drosten and M. Müller) were seeded at 2×10^6^ cells in a 75 cm^2^ flask in Dulbecco’s modified Eagle’s medium (DMEM, supplemented with 10 % (v/v) fetal calf serum (FCS), 1 % (v/v) non-essential amino acids, 100 IU/mL penicillin, 100 μg/mL streptomycin and 2 mM L-Glutamine). After 24 h the cells were inoculated with 100 μl of either wild type virus hCoV-19/Germany/BY-Bochum-1/2020 (GISAID accession ID: EPI_ISL_1118929), VOC B.1.1.7_RKI-0026_B.1.1.7 (GISAID accession ID: EPI_ISL_751799) or the VOC B.1351 RKI-0029_B.1.351 (GISAID accession ID: EPI_ISL_803957). Spike domains of strains were checked for lineage features prior to assays in the context of routine diagnostics (primer kindly provided by René Scholtysik, University Hospital Essen; details about sequences and cycling conditions available upon request). Three days post infection and upon visible cytopathic effects virus suspension was harvested by collecting the supernatant and subsequent centrifugation for 5 min at 1,500 rpm to remove any cell debris. The virus suspensions were aliquoted and stored at −80 °C until further usage.

### Carrier assay

To analyze viral stability on different surfaces we performed time kinetics and studied viral stability over 48 h. Therefore, stainless steel disk, disks sputtered with copper or silver, the inner layer of surgical masks and Filtering Face Piece 2 (FFP2) masks were inoculated with 5 × 10 μL of test virus suspension. The test suspension contained 9-parts virus and 1-part interfering substance (bovine serum albumin [BSA], 0.3g/L in phosphate buffered saline [PBS] according to EN 5.2.2.8) and was adjusted to 5×10^6^ TCID_50_/mL. Immediately, 10 min, 1 h, 24 h and 48 h after virus inoculation on the different surfaces they were placed aseptically in a 2 ml DMEM (without FCS) harboring container and vortexed for 60 s. To determine the amount of recovered infectious virus from the test specimen an end-point-dilution assay was performed on Vero E6 cells to calculate the remaining TCID_50_ according to Spearman and Kärber [6, 7].

### Quantitative suspension assay

To test susceptibility to disinfection, viruses were exposed to 20, 30, 40, 60 and 80 % (v/v) ethanol for 30 s or to hand soap (Lifosan® soft, B. Braun Medical AG, diluted 1:49 in water) for 30 s, 1 min, 5 min and 10 min. Therefore, 8-parts ethanol or hand soap were mixed with 1-part interfering substance (BSA, 0.3g/L in PBS according to EN 5.2.2.8) and 1-part virus adjusted to 5×10^6^ TCID_50_/mL. The suspensions were incubated for the indicated time periods and residual viral infectivity was determined by performing an end point dilution assay on Vero E6 cells.

### Heat inactivation

To access susceptibility towards heat virus suspension were incubated for 1 min, 5 min, 10 min and 30 min at 56 °C. Thus, 9 parts virus adjusted to 5×10^6^ TCID_50_/mL were mixed with 1 part interfering substance (BSA, 0.3g/L in PBS according to EN 5.2.2.8) and incubated for the indicated time periods. Reduction of viral titers were examined by end point dilution assay to calculate TCID_50_ values.

## Results

In order to address if the newly emerged VOC B.1.1.7 and B.1.351 were equally susceptible towards different inactivation strategies as the wild type virus we compared viral inactivation upon usage of ethanol, a common ingredient of several disinfectants and recommended by the World Health Organization (WHO) in resource limited countries [8]. Viruses were exposed towards increasing concentrations of ethanol for 30 s and residual viral infectivity was determined by endpoint titration. In accordance to previous results, all three viral variants could be efficiently inactivated upon treatment with at least 30 % (v/v) ethanol for 30 s, confirming equal susceptibility towards disinfection (**Figure 1**). Since disinfection procedures are mainly recommended in clinical setups, we next addressed the virucidal activity of conventional hand soap. SARS-CoV-2 variants were inoculated with a 1:49 dilution of commercially available hand soap and viral infectivity determined after different time points. All viral variants were effectively inactivated after exposure towards soap within 1 - 5 minutes, supporting current hygiene measures (**Figure 1**). Next, we addressed susceptibility of the three strains towards heat (56°C) and observed a decrease in viral titers towards background levels within 30 min. Importantly, inactivation kinetics were comparable between all viral variants (**Figure 1**). Although SARS-CoV-2 is mainly transmitted through respiratory droplets and aerosols exhaled from infected individuals transmission via fomites cannot be excluded. Viral stability was examined on representative materials surfaces: silver, copper and stainless-steel discs for up to 48 h, using an initial virus concentration of 9.2 × 10^6^ TCID_50_/mL. Importantly, all variants remained infectious on the different surfaces for 48 h and compared to the wildtype virus no differences in the relative infectivity were observed (**Figure 2A**). In order to mimic a potential contamination of on protective masks by infected individuals, we contaminated the inside of either a surgical mask or a FFP2 mask and analyzed viral stability for all variants. Again, comparable residual titers of all VOCs were observed over time (**Figure 2B**). In conclusion, the currently circulating VOC did not exhibit enhanced surface stability or differences in disinfection profiles indicating that current hygiene measures are sufficient and appropriate.

**Figure 1:**
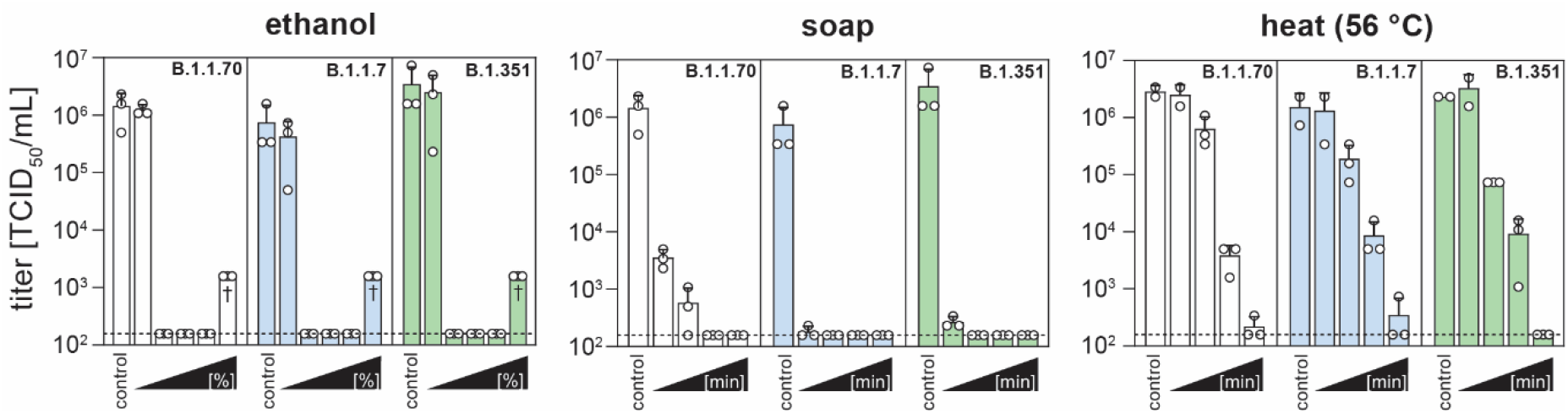
Inactivation of SARS-CoV-2 B.1.1.7 and B.1.351 variants compared to B.1.1.70 (wild type). Residual titer (TCID_50_/mL) of B.1.1.70 (white bars) B.1.1.7 (blue bars) and B.1.351 (green bars) variants after inactivation via heat (56 °C, left panel) for 1, 5, 10 and 30 min (left to right), soap (middle panel) for 30 s, 1, 5 and 10 min (left to right) and ethanol (right panel, 20%, 30%, 40%, 60% and 80%, left to right). Depicted are the individual replicates as dots and the mean as bars ± SD; dashed line indicates lower limit of quantification (LLOQ) of the limiting dilution assay. † denotes elevated LLOQ due to cytotoxicity.

**Figure 2:**
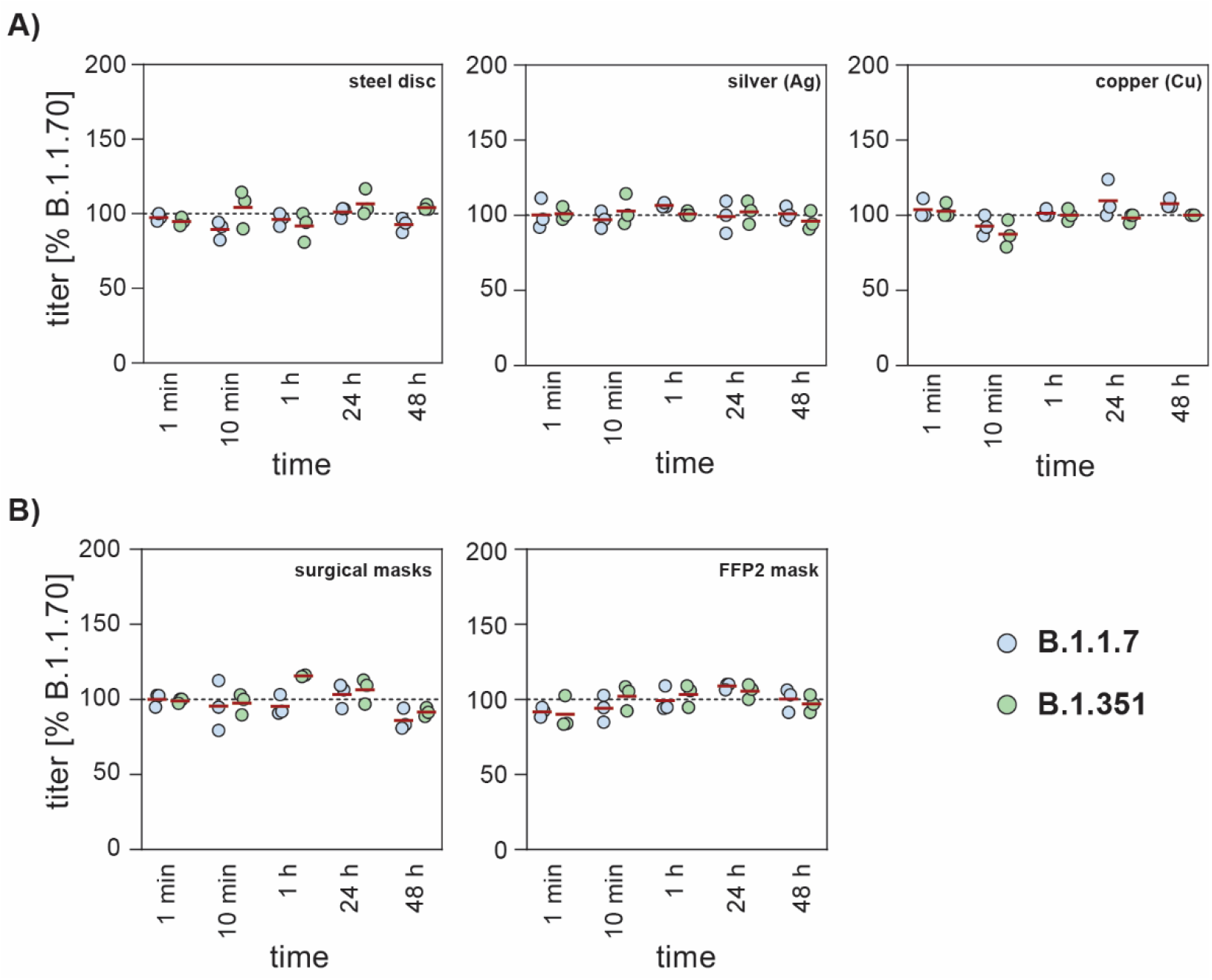
Relative stability of SARS-CoV-2 B.1.1.7 and B.1.351 variants to B.1.1.70 (wildtype). SARS-CoV-2 stock solutions were applied on different surfaces and recovered after the indicated times and residual titer was assessed via limiting dilution assay (TCID_50_/mL). Normalized stability of B.1.1.7 (blue dots) and B.1.351 (green dots) variants on A) stainless steel discs and disks sputtered with copper or silver and B) on the inner layer of surgical masks and Filtering Face Piece 2 (FFP2) masks relative to wild type (dashed line). Depicted are the individual replicates as dots and the mean as red lines.

## Discussion

The currently circulating VOCs, including B.1.1.7 and B.1.351 have shown a strong increase in incidences in various countries. In particular, the B.1.1.7 strain has been suspected to display a 43–90% higher reproduction number compared to preexisting variants [3, 9]. However, the exact mechanisms underlying the increased transmission rates are still under investigation. Given the challenges during the rollout of COVID-19 vaccines, current prevention measures are based on the “swiss cheese model” [10], including a combination of different intervention strategies. In most countries, physical distancing, face covers and hygiene measures are the main strategies to lower virus spread. Therefore, it is essential to address if current hygiene strategies are sufficient and appropriate to prevent transmission of newly emerging VOCs. Especially in the hospital setting, viral disinfection is crucial given the large number of infected patients with high viral loads in a limited space. Several disinfectants are based on ethanol which has been shown to efficiently inactivate CoVs within a very short time frame [11]. In agreement with this, we observed a comparable susceptibility of all viral variants tested towards a minimum of 30 % ethanol upon 30 s exposure, indicative of similar disinfection properties. Since disinfections are not recommended for the daily use, we further examined the virucidal efficiency of common household soap. Soaps contain a mixture of surfactants, which can act directly antiviral upon insertion into the lipid envelope thereby leading to the disintegration of the virus within minutes [12, 13]. However, given that common day-to-day practices do normally not include soaping of hands for several minutes, additional effects can include viral elution from the hand surface due to the adsorptive properties of soap that results upon hand rubbing and subsequent washing in successful removal of the viral particles [14]. We observed an efficient inactivation of all variants within 30 s exposure and upon 5 min all viral variants were completely inactivated. Of note, contact times can differ depending on the ratio of soap and water. Interestingly, we observed slight differences with a minimal residual infectivity after 30 s and 1 min for the wildtype in contrast to the tested VOCs. However, these could be attributed to a variety of factors and do not necessarily reflect changed biological properties of the viruses. In order to minimize the risk of SARS-CoV-2 transmission while handling and processing of clinical specimens, standard precautions involve different inactivation procedures to reduce or abolish infectivity. Heat inactivation protocols are commonly used for a variety of subsequent applications, therefore, we aimed to address the susceptibility of VOCs towards treatment with 56 °C for different times. As described before, a 30 min treatment with 56 °C is sufficient to efficiently abolish infectivity, with no differences between the VOCs. Transmission via contaminated surfaces (fomites) is not considered to be a main route of infection, nevertheless given the high transmission rates questions regarding changed environmental stability were being raised. Surface stability for several days has been described under laboratory conditions for several coronaviruses [15–17]. Using different surfaces, we did not observe any differences regarding viral decay kinetics. Importantly, we observed prolonged stability of all variants on face masks, highlighting the importance of exchanging masks regularly and the risk of shared masks. Of note, in contrast to other publications [18], we did not observe an antiviral effect of silver surfaces on SARS-CoV-2. This is in contrast to copper, for which antiviral properties have been described before and could be confirmed in this study [19]. In conclusion, our results suggest that current hygiene measures are appropriate and effective against the currently circulating VOCs.

## Acknowledgements

We would like to thank all members of the Department for Molecular & Medical Virology for helpful suggestions and discussions.

